# OPTIKA, a new high content kill-kinetic assay to longitudinally assess *in vitro* drug combinations against *Mycobacterium tuberculosis*

**DOI:** 10.64898/2026.05.10.724062

**Authors:** María Pilar Arenaz-Callao, Pablo Gamallo, Alfonso Mendoza-Losana, Santiago Ferrer-Bazaga, Rubén González del Río, Santiago Ramón-García

**Author notes:** Corresponding author. Mailing address: Department of Microbiology, Pediatrics, Radiology and Public Health. Faculty of Medicine. University of Zaragoza. C/ Domingo Miral s/n. 50009. Zaragoza, Spain. Current addresses: Global Health Medicines R&D, GSK, Tres Cantos, Madrid, Spain. Departamento de Bioingeniería, Universidad Carlos III de Madrid (UC3M), 28903 Madrid, Spain.

## Abstract

*In vitro* methods to characterize drug combinations typically involve phenotypic screenings using checkerboard assays (CBA) or, more recently, DiaMOND. Such approaches rely on the Fractional Inhibitory Concentration Index (FICI), a fixed-time measurement of growth inhibition that, nonetheless, necessitates secondary validation by time-kill assays (TKA). Longitudinal time-kinetics of bacterial killing are considered the gold standard *in vitro* proxy for antimicrobial activity, but they required increased assay complexity, particularly against the slow growing *Mycobacterium tuberculosis*.

Here, we developed a new methodology named OPTIKA (Optimized Time Kill Assays) that enhances the capacity of traditional TKA by over 1000-fold. This allows for easy and dynamic examination of *n*-way drug interactions by simultaneously monitoring bactericidal and sterilizing capacities in a longitudinal manner. We then replicated previous DiaMOND studies and performed comparisons using CBA and OPTIKA methodologies. We demonstrate that selection of the efficacy parameters (either routed on bacteriostatic, bactericidal or sterilizing properties) affects the interpretation of *in vitro* drug interactions and, consequently, its potential translational value. The increased assay throughput provided by OPTIKA offers a novel framework for developing tuberculosis treatment regimens.

**Teaser:** OPTIKA is a new methodology that increases time-kill assay performance against *Mycobacterium tuberculosis* by over 1,000-fold

## INTRODUCTION

Tuberculosis (TB) remains a leading cause of mortality worldwide due to an infectious agent, *Mycobacterium tuberculosis* (*Mtb*). The global rise in numbers of people falling ill with TB that started during the COVID-19 pandemic has slowed and are stabilizing. According to the 2025 WHO global TB report, an estimate of 10.7 million individuals contracted TB and 1.23 million fatalities were recorded in 2024. Of those, drug-resistant (DR) TB impact on nearly four hundred thousand people around the world, resulting in a public health crisis and representing a health security threat in several countries (*1*).

The current standard 6-months treatment for drug susceptible (DS)-TB was developed decades ago; it involves an initial two-month intensive phase of isoniazid, rifampicin, pyrazinamide and ethambutol followed by a four-month dual therapy of isoniazid and rifampicin (*2*). In 1950, shortly after the discovery of *p*-aminosalicylic acid and streptomycin, a clinical study demonstrated that a combination of both drugs was more effective and reduced acquired drug resistance. In 1952, isoniazid was discovered and quickly assessed in numerous clinical trials as a combination therapy with available anti-TB drugs. The triple therapy consisting of isoniazid and *p*-aminosalicylic acid over the course of 18 to 24 months, alongside pyrazinamide for the initial six months, resulted in predictable recoveries for 90-95% of patients. As such, it became the standard treatment for TB for the ensuing 15 years. In the 1960s, the use of ethambutol instead of *p*-aminosalicylic acid led to a regimen lasting 18-months that was better tolerated. In the 1970s, rifampicin was included with streptomycin, isoniazid and ethambutol, reducing the treatment time to 9-12 months. Finally, in the 1980s, pyrazinamide replaced streptomycin (*2–4*) and the treatment duration was then reduced to six months, resulting in the current standard anti-TB regimen, that has remained unchanged for decades. When followed correctly, the success rate is high; however, in spite of being a lengthy treatment that can cause liver damage, proper compliance requires patients to take an average of ten pills per day in the intensive phase, which is poorly tolerated by a significant number of patients (*5*). This results in low adherence and the emergence of drug-resistant tuberculosis (DR)-TB, a form of the disease with more limited treatment options. Recently, a 4-month regimen composed of rifapentine, isoniazid, pyrazinamide and moxifloxacin for the treatment of DS-TB was recommended by the WHO, although its implementation at country level is proven challenging due to rifapentine shortages and the concern of emergence of fluoroquinolone resistance in certain parts of the world (*6*).

Despite this achieved progress, there is a continuous need to identify new anti-tubercular drugs and regimens to counteract the rapid emergence of drug resistance affecting the newest regimens and to develop shorter therapies (*7, 8*). Recent decades have seen the revitalization of the antitubercular pipeline (*9*), with several drugs in different clinical phases, some of which being new chemical entities (*10*). Understanding how to optimally combine these compounds into new therapeutically effective multi-drug regimens remains an unresolved question. Since it is not feasible to test all options randomly in the clinic, it is critical to identify ways to prioritize the most promising combinations early in pre-clinical development.

Traditional *in vitro* methods to assess new potential drug combinations involve empirical phenotypic screens for *in vitro* drug interactions, specifically looking for synergies, at the microbiological level. This has been usually studied by checkerboard assays (CBA), which can interrogate 2-way drug interactions, but with limited ability to detect 3-way or higher order drug interactions; CBA are resource intensive and have limited performance capacity. M.C. Berenbaum described a method to test for synergy with any number of drugs (*11*), which was the basis for the implementation performed by Cokol *et al.* (*12*) to evaluate anti-TB drug combinations. This methodology known as DiaMOND (diagonal measurement of *n*-way drug interactions) has significantly streamlined the process of assessing combinations of three or more drugs against *Mtb* (*13*). This is accomplished by reducing the number of interactions that need to be tested to just the most informative wells in the equipotent diagonal of the interaction, thereby increasing throughput capacity (*12*). However, there are some differences between CBA and DiaMOND methodologies: (*i*) *Parameters used for drug interaction calculations.* Basically, in the CBA, FICI values (FICI_90_) are calculated based on concentrations defined by absolute MIC values (which can be assumed as an IC_90_), while DiaMOND performs monotonic adjustments to dose response curves to calculate IC_50_ values and, thus, infer FICI_50_ as the main efficacy parameter; (*ii*) *Cut-off classification of drug interactions are based on FICI values and interactions are classified according to these values.* On the one hand, CBA uses the full two-dimensional interaction space and classifies drug interactions as synergy (FICI_90_ ≤ 0.5), no interaction (FICI_90_ = 0.5-4) and antagonism (FICI_90_ > 4) (*14*), which are considered practical biological limits. On the other hand, DiaMOND simplifies the CBA and only uses dose responses of the drugs alone and the equipotent mixture of the two compounds. DiaMOND relies on the Loewe’s additivity model, in which deviations from FICI = 1 indicate interaction (i.e., FICI > 1 negative and FICI < 1 positive interaction) and incorporates experimental variability, thus classifying drug interaction profiles according to FICI_50_ < 0.85 (synergy), FICI_50_ = 0.85-1.1 (no interaction) and FICI_50_ > 1.1 (antagonism) (*12*). These discrepancies between CBA and DiaMOND classifications of drug interactions exemplify the difficulties on how to define synergy and achieve a standardize criteria (**Figure S1**). Although several mathematical models have been extensively discussed on how accurately define synergy (*15–17*), the translational value of *in vitro* synergy outputs in predicting clinical outcomes of drug combination therapy is still a matter of debate (*18*). One explanation for the observed discrepancies may be that both CBA and DiaMOND are single time-point assays that are inherently based on the use of a fixed-time growth inhibition, the Fractional Inhibitory Concentration Index (FICI), as the measure of dug effectiveness, rather than longitudinal bacterial killing kinetics (*19*).

Time-Kill Assays (TKA) are the most valuable tool for static *in vitro* pharmacokinetic (PK) and pharmacodynamic (PD) studies that rely on enumeration of Colony Forming Units (CFU) at different time points (instead of growth inhibition measurements at a fixed time-point as in the case of CBA, DiaMOND or any other commonly used *in vitro* assay). In addition, TKA serve as the foundation of mathematical modeling of antimicrobial drug activity, being CFU counts the gold standard of bacterial burden used across historical data to measure and compare new drug performance in clinical development programs (*20–22*). However, in TB research, performing TKA is cumbersome due to the slow bacterial growth rate (taking *ca.* 2-4 weeks to form visible colonies) and the need of working in Biosafety Level 3 (BSL3) laboratories. Although recent improved methodologies allow for higher throughput in TKA (*23, 24*), these have a limited throughput capacity and restrict validation of interactions among more than three drugs due to the numerous possible combinations.

Here, we present a new methodology called OPTIKA (Optimized Time Kill Assays), which substantially enhances the capacity of conventional TKA by: (*i*) enabling cost-effective, longitudinal and dynamic investigation of *n*-way drug interactions through a CFU-free methodology performed on user-friendly 96-well plate format; (*ii*) increasing throughput by over 1000-fold and; (*iii*) decreasing readout times by over two weeks. OPTIKA provides a direct measurement of effect on bactericidal activity and, more importantly, sterilizing capacity, which is expected to represent a more relevant translational *in vitro* surrogate for clinical effectiveness.

## RESULTS

### Drug interaction profiles are dependent on methodology and parameters used for their classification

To better understand whether differences between CBA and DiaMOND methodologies could affect the interpretation of drug interactions against *Mtb*, combinations tested by Cokol *et al.* (*12*) (from now on DiaMOND_Cokol_) were replicated in an 8×8 checkerboard format in 96-well plates combining the nine anti-TB compounds in all possible 36 pairwise combinations. DiaMOND_Cokol_ data were compared to our own experimentally generated data and interactions classified according to each methodology. First, we compared how these 36 pairwise combinations classified according to DiaMOND_Cokol_ (i.e., based on FICI_50_) correlated with our CBA data (i.e., based on FICI_90_). The comparison showed a low correlation, with only 8 out of the 36 combinations providing a matching interaction profile. We also found an overestimation of antagonistic interactions by the DiaMOND methodology versus no interaction profiles reported by the CBA methodology (**Figure 1A**). Next, we selected the equipotent diagonal of the interaction in our data (DiaMOND_own_) and analyze it according to the DiaMOND methodology. For this, first, we evaluate the impact of the model used for interaction calculations: in dose response studies, IC_50_ values are typically calculated by fitting a four-parameter logistic regression model to the data points; however, in the DiaMOND methodology IC_50_ values are calculated by a monotonical decreasing model of the experimental data. We thus confirmed that the model used to fit the dose response data had no major implications on the classification of the combinations (**Figure S2**). We then analyzed our own data according to the DiaMOND methodology (i.e., FICI_50_, monotonical decrease) and compared it to the DiaMOND_Cokol_ data: although the observed correlation remained poor, with only 17 out of the 36 combinations, a slight improvement was observed, stressing the importance of the parameter used for analysis (i.e., FICI_50_ *vs.* FICI_90_) (**Figure 1B).** Calculations based on IC_50_ values consistently delivered higher FICI values irrespectively of the methodology used (i.e., CBA or DiaMOND) compared to those obtained based on IC_90_ values; thus, overestimating the number of antagonistic interactions, especially when applying more restrictive cut-offs. Our data indicated that the inhibitory parameter used (IC_50_ *vs.* IC_90_) did have a significant impact on such classification of drug interactions (**Figure S3**). Introducing a bactericidal readout (Fractional Bactericidal Concentration Index, FBCI_99.9_) showed that bacteriostatic profiles were the dominant trend in this combination set (23 out of the 36 combinations were bacteriostatic) with only 4 out of 13 remaining combinations matching DiaMOND_Cokol_ classification and showing a synergistic bactericidal interaction (**Figure 1C**).

**Fig. 1.**
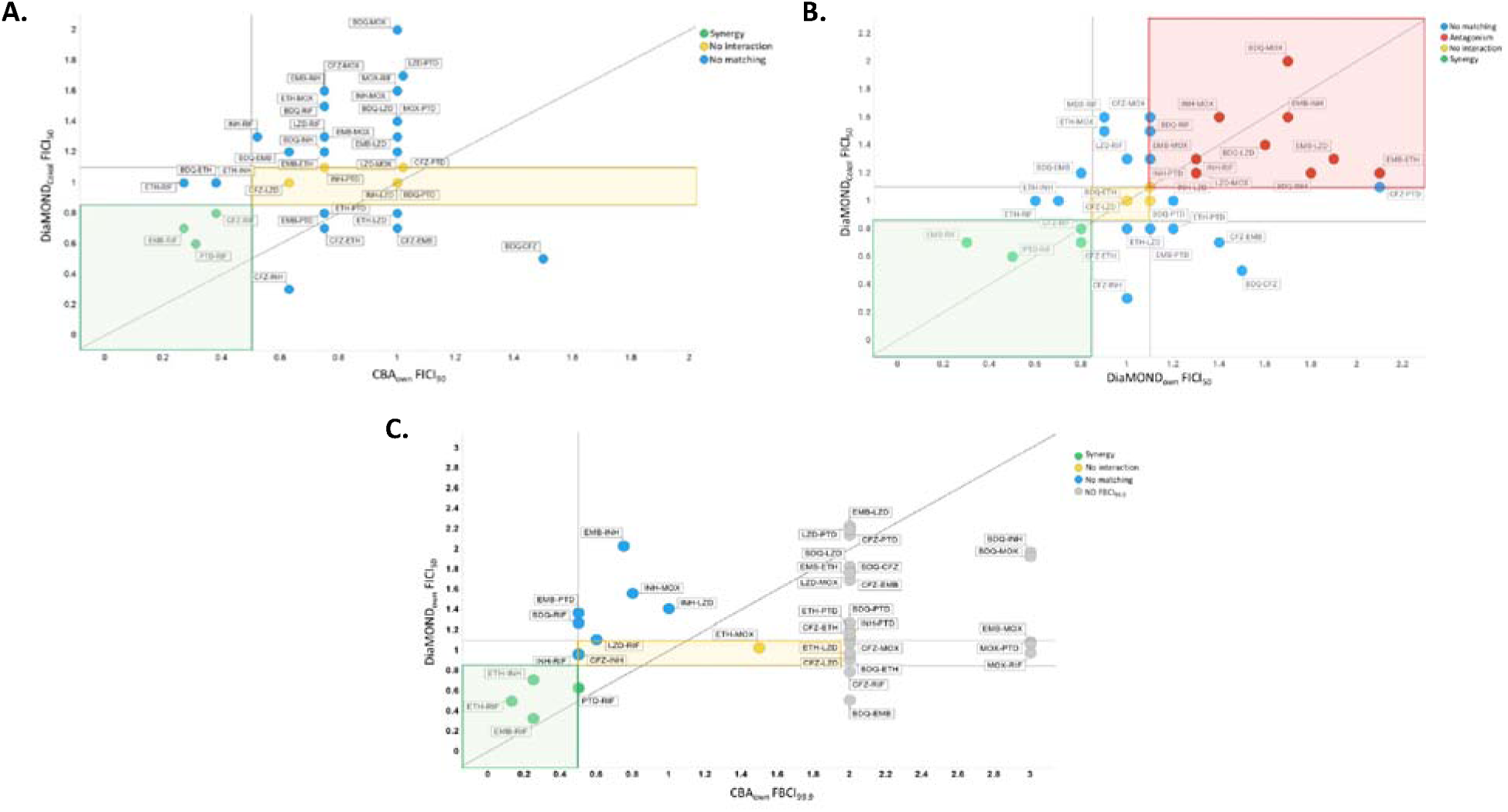
Drug interaction classifications according to DiaMOND and CBA methodologies. **(A)** DiaMOND data by Cokol *et al.* (*12*) reported as FICI_50_ (DiaMOND_Cokol_ FICI_50_) compared to internally generated data analyzed according the CBA methodology and reported as FICI_90_ (CBA_own_ FICI_90_). Correlation of 8 out of 36 combinations (8/36) tested **(B)** DiaMOND_Cokol_ FICI_50_ compared to internally generated data analyzed according to the DiaMOND methodology (DiaMOND_own_ FICI_50_). Correlation of 17 out of 36 combinations (17/36) tested. **(C)** Internally generated data analyzed by DiaMOND reported as FICI_50_ (DiaMOND_own_ FICI_50_) compared to internally generated data reported as FBCI_99.9_, analyzed according to the CBA methodology (CBA_own_ FICI_99.9_). Twenty-three out of the 36 pair-wise combos (23/36) were bacteriostatic. Of the 13 combinations showing a bactericidal effect, only 4 (4/13) correlated with Cokol *et al.* (*12*). Circle legends: Green, synergy; Yellow, no interaction; Blue, no matching combinations; Grey, non-determined. Background zone legends (according to FICI cut-offs): Green, area for synergy combinations classified by both DiaMOND and CBA; Yellow, area for no interaction combinations; White: area for combinations classified differently by both methods. Straight dotted line: perfect correlation zone (x=y). FICI, fractional inhibitory concentration index; CBA, checkerboard assay; DiaMOND, diagonal measurement of *n*-way drug interactions. Bedaquiline (BDQ), clofazimine (CFZ), ethambutol (EMB), ethionamide (ETH), isoniazid (INH), linezolid (LZD), moxifloxacin (MOX), pretomanid (PTD), rifampicin (RIF).

### OPTIKA, a new high content longitudinal kill-kinetic methodology to assess drug combinations

To overcome limitations of growth inhibition-based assays, we developed a new methodology named OPTIKA (Optimized Time Kill Assays) that increases the capacity of traditional TKA by more than 1,000-fold (**Table S1**). OPTIKA is based on the “charcoal agar resazurin assay” (CARA) (*25*), which replaces the use of CFU by a resazurin-based fluorescence readout (**Figure 2**). In brief, *Mtb* is exposed to different drug combinations in 96-well plates and, at every time point, a small volume is spotted onto CARA plates. In parallel, a calibration curve is built with 10-fold serial dilutions of an untreated exponentially growing *Mtb* culture with a known cell density. CARA plates are revealed with resazurin. The fluorescence linear range of the calibration curves is fitted to a linear regression so that fluorescence of treated samples could be converted to log_10_ CFU/mL using the calibration curve equation at the corresponding time point.

**Fig. 2.**
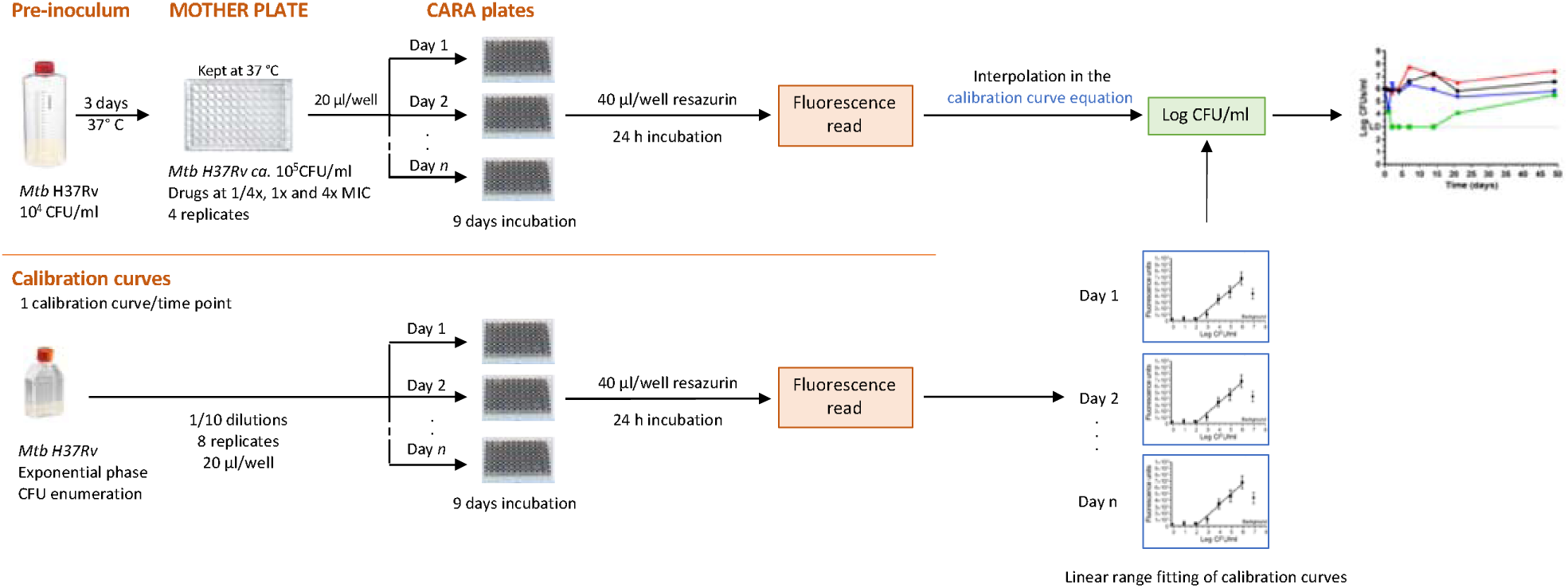
OPTIKA protocol schema. Mother plates with appropriate drug combinations previously dispensed are inoculated with a 3-day old culture of *M. tuberculosis*. At designated time points, aliquots from mother plates are transferred into empty CARA plates. In parallel, a culture with known bacterial cell concentrations is used to generated a calibration curve at every time point. After 9 days of incubation, CARA plates are read following addition of resazurin. The fluorescence linear range of the calibration curves is fitted to a linear regression so that fluorescence of treated samples is converted to Log_10_ CFU/mL using the calibration curve at the corresponding time point.

Head-to-head comparison of OPTIKA with traditional CFU-based killing curves showed resazurin-based OPTIKA reproducing CFU-based readouts (**Figure S4**).

### Diverse kill kinetic profiles are successfully identified with OPTIKA

The OPTIKA methodology allows for the generation of a large amount of time-kill kinetic data of both combinations and corresponding single drugs, enabling the identification of several killing profiles (**Figure 3A**). For example: (*i*) the combination of rifampicin and isoniazid showed a fast-killing profile with the combination killing faster than any of the single drugs alone, as shown by the slope at day 2. The low bacterial load reached by the combination was maintained until day 14. However, this was followed by a rapid rebound starting on day 14 and no differences were observed compared to controls at endpoint (day 49); (*ii*) the previously described synergistic combination of ethambutol and rifampicin (*26*) was confirmed here by OPTIKA. The limit of detection of bacterial killing was rapidly reached by the combination without resumption of bacterial growth. In contrast, they showed different degrees of bacterial killing when single drugs were assayed separately, although without reaching the limit of detection at any sampling time. Worth noting that this strong interaction was maintained up to 49 days, despite rifampicin having a half-life in the assay media of ca. 6 days (*26*); finally, (*iii*) an example of paradoxical antagonism, or slow interaction, was observed when isoniazid was combined with bedaquiline. No positive interaction of the combination was observed until day 7, being the combination less potent than isoniazid alone. However, a bactericidal effect was observed at later time points, being the combination able to prevent the typical development of drug resistance after *in vitro* exposure to isoniazid (*27*). At day 14, the limit of detection was reached and no detectable bacteria were observed over the time course, with the final outcome classified as favorable interaction according to OPTIKA.

**Fig. 3.**
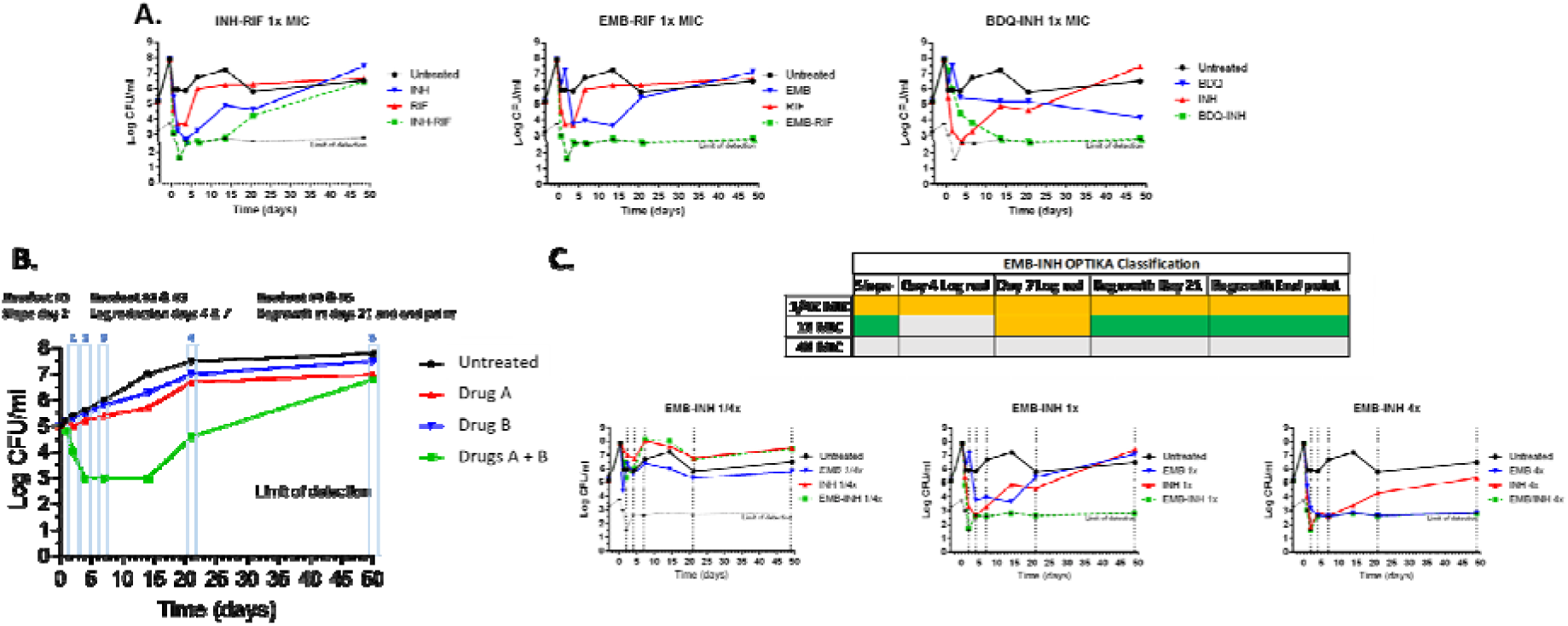
OPTIKA killing profiles and parameterization. **(A)** Different combinations showed various interaction profiles. **(B)** Drug interactions were determined based on five parameters of the time-kill curve: slope at day 2, log reduction at days 4 and 7, and bacterial regrowth at days 21 and 49. **(C)** The killing profile of the EMB-INH combinations is compared against the killing profile of the related single drugs. Green: favorable interaction, i.e., >2 Log_10_CFU/mL reduction in the combination compared to the most active compound alone. Orange: no interaction, i.e., <2 Log_10_CFU/mL reduction in the combination compared to the most active compound alone. ND: non determined, i.e., when the activity of one of the compounds alone reaches the limit of detection, thus masking any potential drug interaction. Red (not shown): non-favorable interaction, i.e., >2 Log_10_CFU/mL increase in the combination compared to the most active compound alone. BDQ, bedaquiline; EMB, ethambutol; INH, isoniazid; RIF, rifampicin. Drugs are tested according their respective MIC values.

In order to simplify the representation of time-kill kinetic data, five parameters were considered for combination classification by OPTIKA: initial bacterial killing rate or slope, bacterial burden reduction at both days 4 and 7, and the ability of drugs to prevent growth of the bacteria over time, with readout at both day 21 and endpoint (day 49) (**Figure 3B**). These parameters where thus used to classify the combinations based on the interaction profile, i.e., favorable interaction, no interaction, non-favorable interaction or not determined (when one of the drugs alone exerts all the anti-bacterial activity by itself, thus masking the effect of any potential interaction in combination). The combination of ethambutol plus isoniazid was used to exemplify these calculations (**Figure 3C**). At concentrations 1/4xMIC values, none of the five parameters identified any favorable combination. When the concentration was increased at 1xMIC values, the slope and regrowth parameters identified a favorable interaction, being the combination more potent than any of the two drugs alone. However, concentrations of 4xMIC values were unable to define any interaction since both ethambutol and isoniazid alone were active by themselves to the limit of detection at different time points, thus masking the effect of the combination. These results suggested that 1xMIC matching concentration values were the most suitable to identify drug combination interactions.

### OPTIKA allows the dynamic classification of drug combinations based on bactericidal and sterilizing readouts

We performed OPTIKA with the 36 pairwise combinations previously studied (*12*) to evaluate their time component and estimate their bactericidal/sterilizing potencies. The high-throughput achieved by OPTIKA allowed us to test different concentration patterns, thus reducing the number of false positive or negative results. **Figure 4** shows the drug interaction classification according to the parameters previously described for every concentration of the above-mentioned combinations: sub-MIC (1/4xMIC), MIC (1xMIC) and over-MIC (4xMIC), which could be explained considering the dynamic profile of the drug interactions in the time-kill assays. Interactions observed at shorter incubation times may be reversed in the long term or vice-versa. This happened, for example, in the clofazimine-pretomanid (CFZ-PTD) at 1xMIC and bedaquiline-ethambutol (BDQ-EMB) at 1xMIC combinations. Considering just one parameter, combinations could display different interactions profiles depending on the concentration tested. For example, the slope of moxifloxacin-isoniazid (MOX-INH) combination showed different classifications at sub-MIC, MIC or over-MIC. Thus, all indicators taken together allowed us to describe the curve profile without a visual inspection of the actual killing curve plots. In general, only strong interactions were identified at sub-MIC values whereas most of the interactions could not be determined at over-MIC values because single drugs alone showed their maximum effect, therefore masking any dynamic window to observe the effect of the combination. In the search of a single parameter to classify the combinations and aiming to overcome limitations of standard methods, slope at day 2 (proxy of bactericidal activity) and long-term incubation were prioritized, being endpoint regrowth (day 49) the chosen parameter as an *in vitro* proxy of sterilization. Again, the 1xMIC condition provided the largest dynamic window and was the preferred concentration to categorize drug interactions, being the most informative for most of the combinations.

**Fig. 4.**
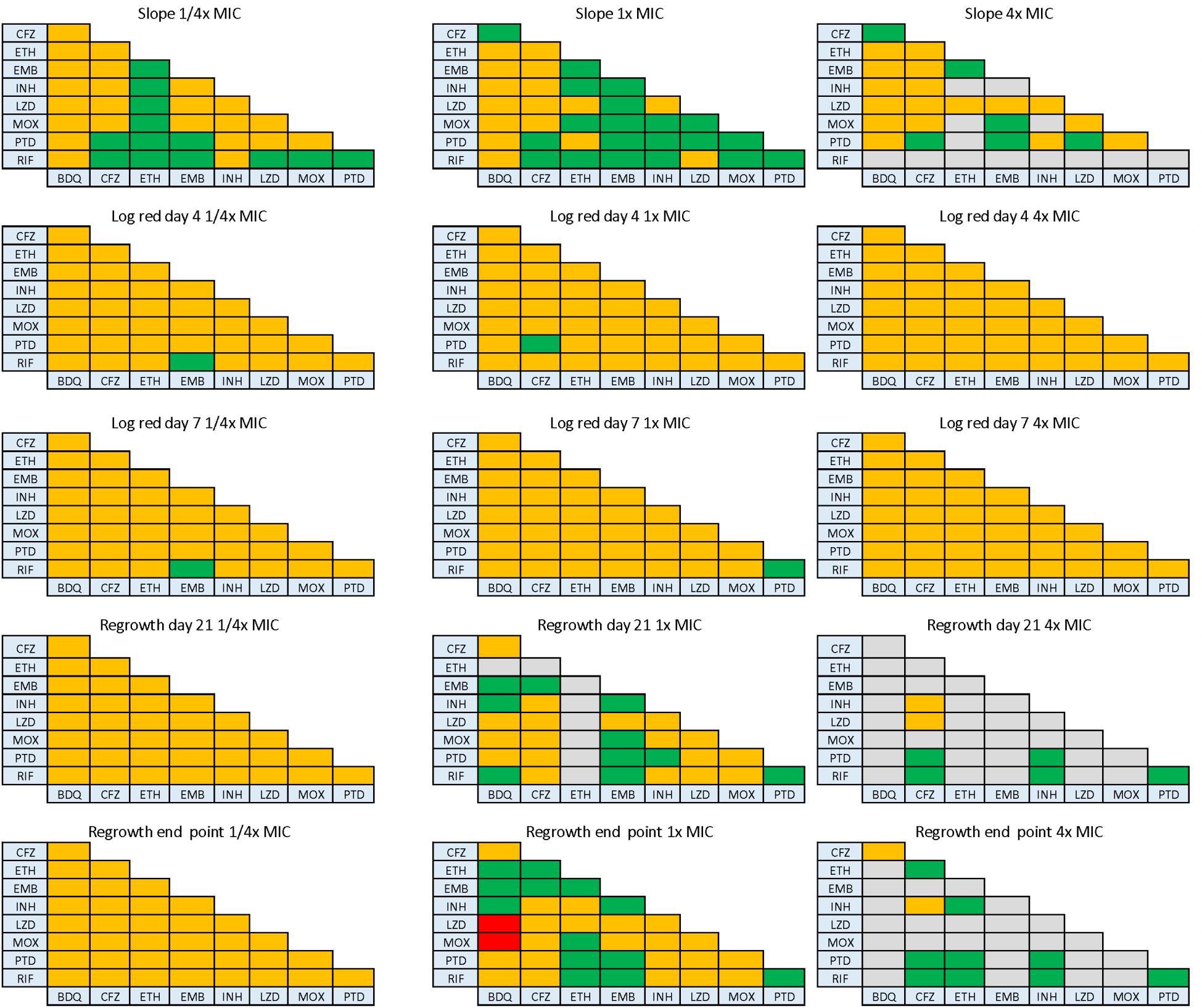
Pair-wise combinations by OPTIKA. The 36 pair-wise combinations described in Cokol *et al.* (*12*) were tested using OPTIKA. Each triangle represents the classification of the set of pair-wise combinations based on one OPTIKA parameter at one determined concentration. Left column: combinations assayed at 1/4xMIC. Middle column: combinations assayed at 1xMIC. Right column: combinations at 4xMIC. Rows represent the different parameters (from top to bottom): slope at day 2, log reduction at day 4, log reduction at day 7, regrowth at day 21 and regrowth at day 49. Green, favorable interaction; Orange, no interaction; Red, non-favorable interaction; ND, interaction not determined due to activity of single drugs to the limit of detection. Bedaquiline (BDQ), clofazimine (CFZ), ethambutol (EMB), ethionamide (ETH), isoniazid (INH), linezolid (LZD), moxifloxacin (MOX), pretomanid (PTD), rifampicin (RIF).

### Bactericidal and sterilizing readouts identified favorable combinations against *Mtb*

The interaction profile of the 36 pairwise reference combinations by the different methods is summarized in **Figure 5A**. Interestingly, all synergistic interactions detected by CBA using the FICI_90_ were classified as favorable interaction by the endpoint regrowth OPTIKA parameter. (bedaquiline-ethionamide, clofazimine-rifampicin, ethambutol-rifampicin, ethionamide-isoniazid, ethionamide-rifampicin and pretomanid-rifampicin). In addition to these six combinations, OPTIKA identified 12 additional favorable pairwise combinations, being four of them (bedaquiline-ethambutol, clofazimine-ethionamide, clofazimine-ethambutol and clofazimine-pretomanid) slow interactions for which favorable effects were only detectable by long term incubation with a similar profile than the one described for bedaquiline-isoniazid (**Figure 3A)**. Two combinations were robustly classified as favorable interaction by all three methods: the well-known favorable combination of rifampicin and ethambutol and rifampicin plus pretomanid (*12*). Data integration confirmed that the use of FICI_50_ (fixed time-point growth inhibition readout) overestimated the presence of antagonistic/non-favorable interactions (as shown by both DiaMOND_Cokol_ data and our DiaMOND_own_ data), while the use of bactericidal readouts (slope day 2) substantially enhanced the potency to identify synergistic/favorable interactions. Regrowth at endpoint was the most restrictive and robust parameter to identify sterilizing combinations (**Figure 5B**).

**Figure 5.**
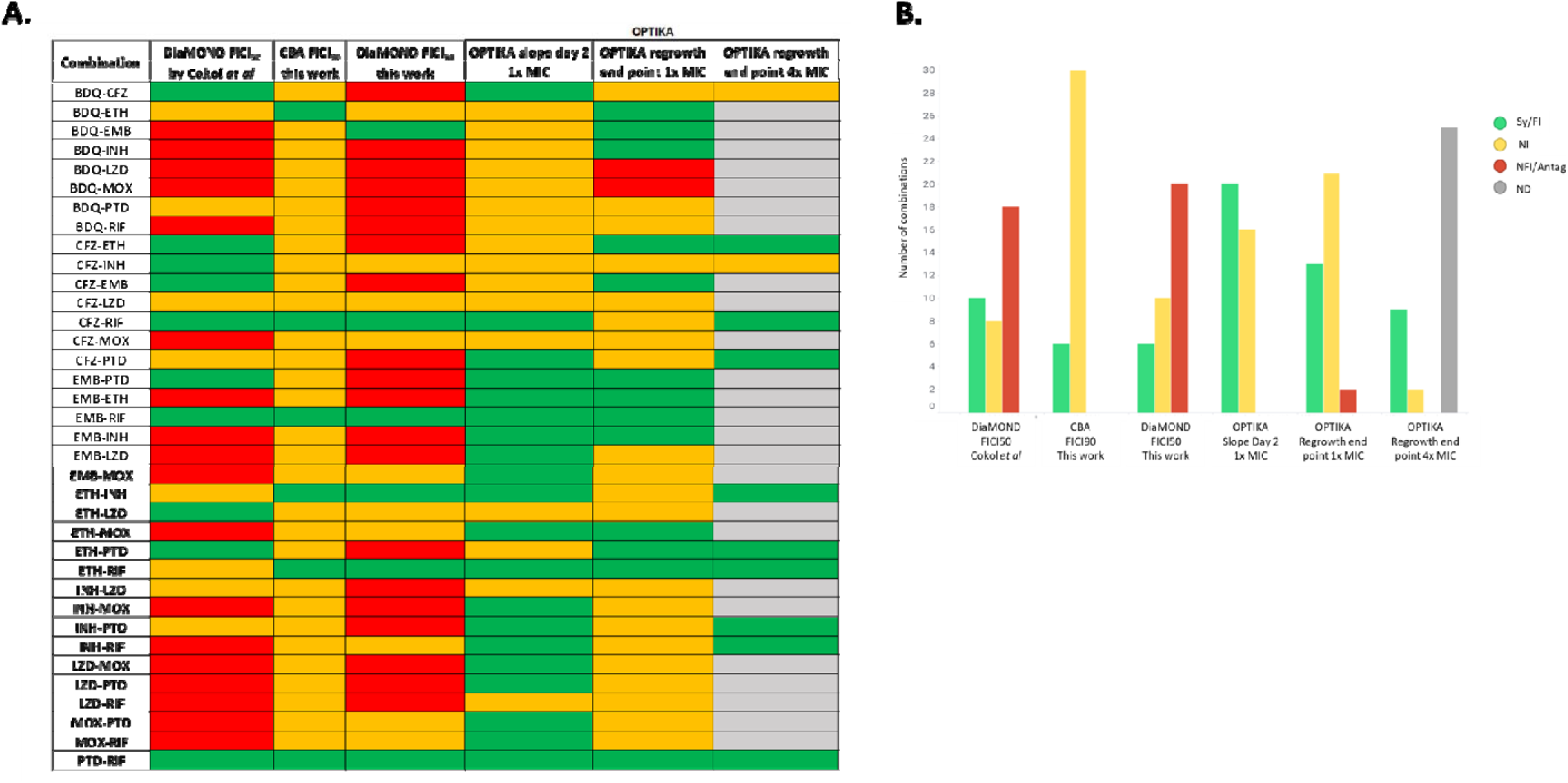
Classification of pair-wise drug interactions according to the different methodologies used in this study. **(A)** The interaction of the 36 pair-wise reference combinations were classified according the three methodologies used in this study: CBA, DiaMOND and OPTIKA and using the appropriate readout parameter, i.e., FICI_50_, FICI_90_ and the OPTIKA parameters “slope day 2” and “regrowth at endpoint” (both at 1xMIX and 4xMIC values). **(B)** Number of combinations classified according to the different methods. Green: synergy/favorable interaction; Orange: no interaction; Red, antagonism/non favorable interaction; Grey: not determined. Bedaquiline (BDQ), clofazimine (CFZ), ethambutol (EMB), ethionamide (ETH), isoniazid (INH), linezolid (LZD), moxifloxacin (MOX), pretomanid (PTD), rifampicin (RIF).

### OPTIKA allows the study of *n*-way drug combinations

Next, we aimed to apply OPTIKA to the study of higher-order combinations. The ten triple drug combinations previously described by Cokol *et al.* (*12*) were tested by OPTIKA at all possible triple combinations of five drugs (bedaquiline, clofazimine, isoniazid, pretomanid and rifampicin). For this, similar to pairwise studies, matching concentrations (i.e., 1/4xMIC, 1xMIC and 4xMIC) of each single drug were combined in the 3-way combos. Nine and eight combinations showed a favorable interaction by OPTIKA according to “slope at day 2” or “regrowth at endpoint” parameters, respectively, in at least one of the three concentrations tested. In contrast, only two combinations were classified as synergistic by DiaMOND_Cokol_ **(Figure 6).** Due to the increased capacity of OPTIKA to perform time-kill assays, when assessing *n*-way combinations it is thus possible to interrogate the contribution of the lower order combinations to the overall performance of the high-order combination, as shown in our deconvolution studies **(Figure 6B** and **figure S5)**. For example, in the combination of bedaquiline with isoniazid and rifampicin (BDQ-INH-RIF) there was a positive drug interaction between bedaquiline and isoniazid from day 14, similar to the profile in the triple combination from day 14 to 49. This favorable outcome of the combination could be attributed to the prevention of the well-known isoniazid-resistant bacteria *in vitro* generation. Another explanation could be that the bacterial metabolism reduced by the effect of bedaquiline (*28*) could potentiate the bactericidal activity of isoniazid. The addition of rifampicin to this slow pairwise interaction caused a favorable faster bactericidal interaction that produced the observed strong killing ability in the triple combination from day 1 to day 14. This first part of the kill curve matched with the corresponding part of the pairwise combination formed by isoniazid plus rifampicin. Finally, the combination of rifampicin with bedaquiline showed a bactericidal positive interaction, but weaker than BDQ-INH, that was not able to prevent bacterial regrowth, thus it did not have sterilizing capacity. In summary, the combination of BDQ-INH-RIF showed a rapid bactericidal activity and sterilizing capacity that could be explained by the contribution of the respective pairwise combination.

**Figure 6.**
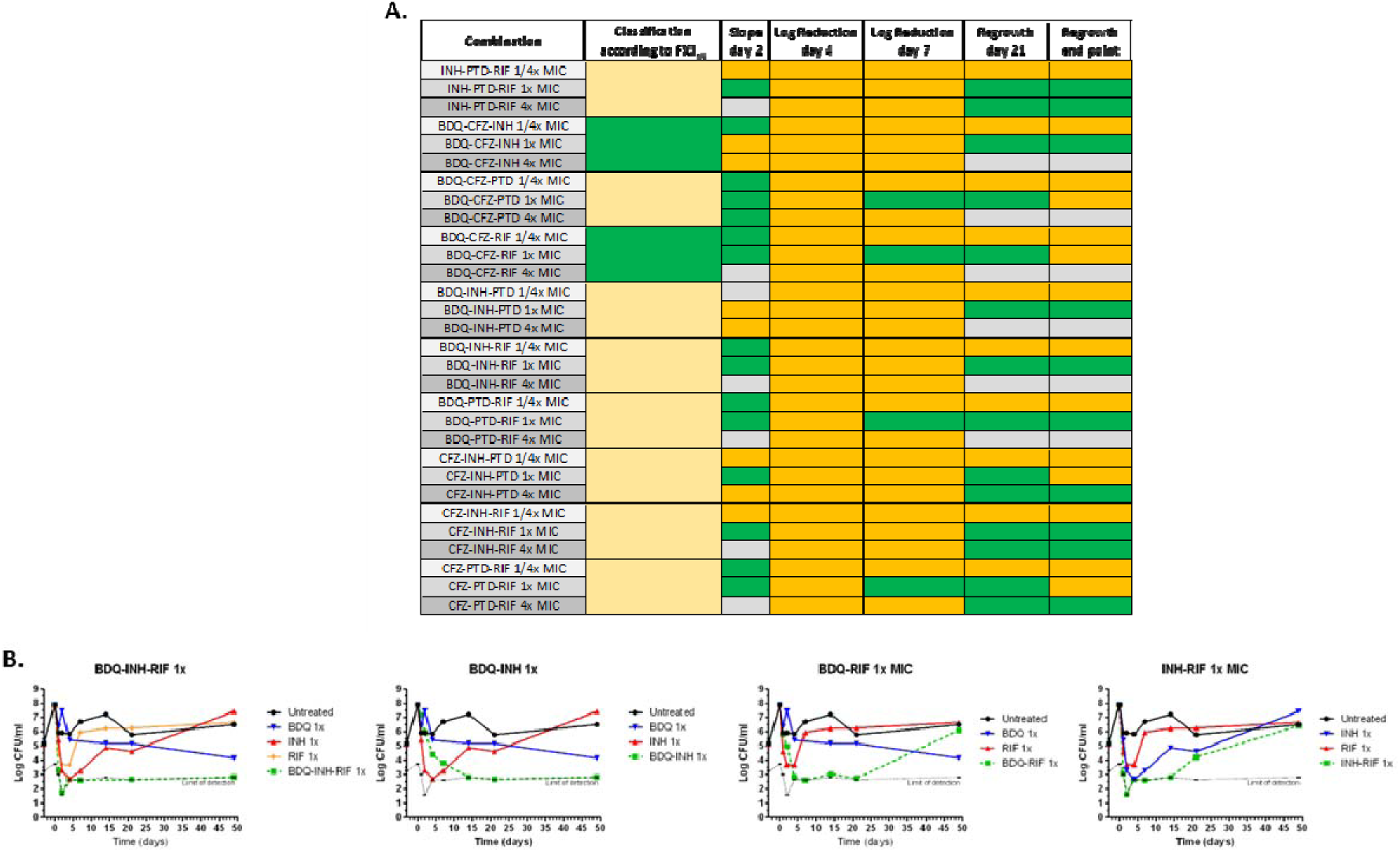
Triple combinations by OPTIKA. **(A)** Classification according to FICI50: data reported in Cokol *et al.* (*12*). Slope day 2: classification according to OPTIKA slope parameter. Log reduction days 4 and 7: classification according to OPTIKA log reduction parameter at days 4 and 7, respectively. Regrowth at day 21 and endpoint: classification according to OPTIKA regrowth parameter at days 21 and 49 respectively. Green: synergy/favorable interaction; Orange/yellow: no interaction; Red (not shown), antagonism/non favorable interaction. Grey: not determined. Bedaquiline (BDQ), clofazimine (CFZ), ethambutol (EMB), ethionamide (ETH), isoniazid (INH), linezolid (LZD), moxifloxacin (MOX), pretomanid (PTD), rifampicin (RIF). **(B)** Deconvolution of the triple combination BDQ-INH-RIF. The killing profile of the triple combination is compared against the killing profile of the related pair-wise combinations. Bedaquiline (BDQ), isoniazid (INH), rifampicin (RIF).

## DISCUSSION

Combination of rapidly acting antitubercular drugs and those with sterilizing capacity are imperative for an effective TB treatment. The development of the 6-month drug regimen (RHZE, rifampicin-isoniazid-ethambutol-pyrazinamide) took approximately 20 years and was solely based on time-consuming trial/error clinical studies (*2*). Recent years have witnessed the emergence of an unprecedent number of new chemical entities with drug-like properties that exhibit anti-mycobacterial activity, with some of these compounds currently undergoing clinical trials (*10*). However, experimental methods for integrating these agents into effective multidrug regimens for TB treatment still remain suboptimal. Recent predictive efforts to inform the design of clinical trials have mostly relied on the integration of data from mouse models of *Mtb* infection and estimates of drug penetration in the caseum of granulomas in rabbit models of *Mtb* infection (*29, 30*). However, animal models require extensive labor-intensive experimentation in costly BSL3 facilities with a significant limitation on the number of potential combinations that can be tested. In addition, the current scientific and regulatory landscape is moving towards the progressive replacement of animal testing with New Approach Methodologies (NAM) (*31, 32*).

*In vitro* assays provide a mean to interrogate a large number of drug combinations by empirical phenotypic screening for *in vitro* synergies at the microbiological level. Traditionally, this was done by the checkerboard assay, which allows bidimensional interrogation of 2-way drug interactions; however, this methodology has a limited power to identify 3-way (or higher order) drug interactions due to the complexity of adding a third *z*-axis to the experimental layout. A new experimental approach was recently developed in the TB field to overcome this limitation by reducing the number of experimental conditions that require testing; this approach, named DiaMOND, vastly simplified the ability to identify favorable interactions of combinations of three, four or even a higher number of drugs (*12, 13*). Both CBA and DiaMOND rely on the calculation of the FICI, an index based on fixed-time-point growth inhibition rather than longitudinal bacterial killing, the latter remaining unmeasured in these assays (*19*). While CBA depend on absolute MIC determinations (i.e., IC_90_ values), DiaMOND mainly uses IC_50_ calculations to define a FICI. In addition, each methodology uses different FICI cut-off values to define the type of drug interaction (*12, 14*), which influence how the interaction of drug combinations are classified and, consequently, could introduce a bias on how certain combinations could be prioritized in the preclinical development process (**Figure S1**).

We replicate the experiments described by Cokol *et al.* (*12*) and found that specific parameters used to describe *in vitro* drug interactions (i.e., IC_50_ and IC_90_, based on growth inhibition, or MBC, based on cidality) do actually influence the interaction criteria of such drug combinations. Moreover, we found that the use of IC_50_ values as an indicator of drug interactions artificially enriched the classification for antagonistic combinations (**Figure 1** and **Figure 5B**), which correlates with previous observations showing that the use of IC_80_ values could favor the identification of synergistic interactions (*33*); the question, then, of whether a mathematical classification should be used to reflect biological significance, remains open. Surprisingly, when we replicated the DiaMOND methodology, we observed that only 17 out of the 36 (17/36) combinations were classified as in Cokol *et al.* (*12*). This almost 50% discrepancy could be attributed to differences in assay conditions: (*i*) Although both laboratories used a *Mtb* H37Rv strain, in the original DiaMOND report, this was a BSL2 pantothenate and leucine auxotrophic strain, while we employed an BSL3 ATCC strain; (*ii*) Due to requirements of the strain, the medium used was also different: 7H9 supplemented with OADC, glycerol, tween, panthotenate and leucine versus a more simple medium composed of 7H9 plus ADC and glycerol, respectively; (*iii*) the assay format was also slightly different: while Cokol *et al.* (*12*) used a 384-well plate format (50 µL, 4·10^6^ CFUs/ml), we employed a 96-well plate format (200 µL, 10^5^ CFUs/ml), which could affect the metabolic state of the bacteria. Finally, (*iv*) activity readout was also different: based on optical density measurements (OD_600_) versus MTT readout, respectively, with slightly better resolution of the MTT readout at detecting bacterial burden (*19*). All these differences, and in the light of recent evidence on the impact of strain diversity (*34*), ask for caution when interpreting and using data generated in a single laboratory. Performing external quality assessments would enhance data confidence and robustness with regulatory value. This is already implemented in the case of well-established methodologies such as the EUCAST MIC determination (*35*), and it has been applied for conventional time-kill assays in a multicenter study where the variability of the technic is robustly assessed against a standardized protocol (*23*).

Combinations identified in growth-inhibition, single time-point FICI-based assays should be validated with informative secondary time-kill assays that are normally based on CFU enumerations, the gold standard of *in vitro* quantification of antimicrobial drug activity. Recently, a novel methodology has introduced the use of large-scale live-cell imaging approaches to quantify bacterial killing in real time at single-cell resolution, demonstrating that lethally - rather than growth inhibition - could be a better proxy of treatment outcomes (*36*). Longitudinal bactericidal assays thus provide essential information that can be integrated into pharmacometrics models to predict Phase IIa early bactericidal assay (EBA) clinical outcomes (*22, 37–39*). However, sophisticated imaging capacities might not be as readily available elsewhere and, despite recent methodological advances in standard TKA (*23, 40*), performing such experiments significantly increases the complexity and duration of *in vitro* combination testing, especially when validating triple or higher *n*-way drug interactions.

To overcome some of these issues, we introduce an *in vitro* methodology that allows for cost-effective high-throughput, facile interrogation of *n*-way drug interactions in longitudinal time-kill assays. This novel technology, called OPTIKA, increases standard TKA capacity by 1,000-fold (**Table S1**) and offers key advantages over conventional growth inhibition approaches for drug combination studies: (*i*) it delivers longitudinal measurements on the activity of drugs alone and in combination, allowing multiple testing of different exposures and combination ratios, and; (*ii*) it quantifies bactericidal and sterilizing capacity of the combinations. OPTIKA also offers substantial improvements over standard TKA: (*i*) it enables high-throughput screening of drug interactions in a convenient a 96-well plate format using a surrogate resazurin readout, as opposed to the laborious CFU-based time-kill assays, and; (*ii*) by combining drugs of interest at sub-inhibitory matching concentrations (i.e., 1/4xMIC or 1xMIC), it enables the identification and characterization of favorable interactions among drug partners. In this study, we have tested up to 3-way drug combinations but, as long as drugs are included at sub-inhibitory matching concentrations, this approach could be easily expanded to study *n-*way drug interactions; the high-throughput capacity of OPTIKA ensures that deconvolution studies of the lower-level drug combinations can be also studied to understand the contribution of each drug to the higher order combination. Finally, we used an experimental design powered to identify favorable interactions (i.e., sub-optimal concentrations); however, by selecting matching over-MIC values (i.e., >4xMIC) antagonisms could be also identified, i.e., drugs alone having full bactericidal and sterilizing capacity, while not combinations.

Due to the large amount of data that OPTIKA can generate, we selected critical parameters as best descriptors of the degree of drug interaction (**Figure 3B**), namely: (*i*) the slope of the killing curve at day 2 and (*ii*) the reduction in Log_10_CFU/mL at days 4 and 7 – as a proxy for bactericidal activity-and; (*iii*) bacterial regrowth-as a proxy for relapse-at days 21 and 49 (endpoint). The ability to longitudinally interrogate with high-content capacity several interaction parameters at different drug exposures constitutes an indubitable added value of OPTIKA (*20*). We were able to maintain *Mtb* cultures in a 96-well plate format for up to 50 days. Such long incubation periods after drug exposure are not common in standard *in vitro* assays, typically lasting from 14 days up to one month. As with any time-kill assay, OPTIKA is a static model that maintains bacteria in the same culture medium, without refreshment, and drugs are only added at the beginning of the experiment without any drug replenishment, which might lead to bacterial drug inactivation or thermal degradation of compounds at 37°C in 7H9 media after the long incubation periods (*41*). Despite this potential lack of selective pressure at later time points, our data showed several combinations with favorable drug interaction at the OPTIKA endpoint (i.e., lack of regrowth compared to drug alone) (**Figures 4-6**). These combinations presented a slow but favorable killing profile including: BDQ-EMB, BDQ-INH, CFZ-ETH, CFZ-EMB and CFZ-PTD, with some of them not being captured by DiaMOND (that relies on early fixed time-point for growth inhibition), emphasizing the importance of long-term sterilizing readouts. Triple combinations with enhanced activity over the lower level pairwise combinations were also identified, such as BDQ-INH-RIF (**Figure 6** and **Figure S5**). This is based on three observations: (*i*) favorable drug combinations can prevent the emergence of drug resistance to individual drugs, such as in the case of INH; (*ii*) in the case drugs in the combination are stable over the incubation period, synergistic interactions are clearly identified, and; (*iii*) in the case, one or more drugs are instable, the OPTIKA endpoint would be detecting the post antibiotic effect (PAE) of drug combinations, likely due to an irreversible bacterial damage. This has been previously observed in other anti-TB drug combinations using traditional CFU-based TKA (*26, 42*). In both scenarios, favorable drug combinations at endpoint are defined as those capable of sterilizing the culture while none of the drugs alone can do so.

In recent years, several translational methods and tools have explored ways to preclinically advance anti-TB compounds and combinations (*22, 39, 43–45*). Translational PK/PD models aim to decrease the rate of attrition when progressing compounds from *in vitro* and *in vivo* assays to clinical stages. They typically determine the human drug exposure that would be required to achieve a therapeutic efficacy equivalent to that obtained in animal models by integrating information from different sources, including *in vitro* data (typically single time-point growth inhibition-based data lacking longitudinal information) (*45*). Their objective is to establish a correlation between dosage and response so as to determine *in silico* ideal treatment regimens in EBA efficacy trials. EBA clinical assays evaluate the bactericidal capacity of a new drug or drug combination over the initial treatment period (typically 2 weeks); however, similar to what we have observed in our TKA experiments (**Figure 5-6** and **Figure S5**), early bactericidal activity does not necessarily correlate with sterilization capacity, which is a *sine qua non* condition for clinical TB cure. Similar observations have been extracted from relapse mouse models, which have high potential to predict treatment shortening and forgiveness of tuberculosis treatment (*46, 47*). The OPTIKA protocol is based in cell lethally (rather than growth inhibition) and it has the capacity to interrogate the sterilization activity of drug combinations at extended time-points. Favorable combinations at endpoint (i.e., no regrowth/sterilization) could be the most informative descriptor to classify *in vitro* drug interactions as a proxy for treatment shortening potential. However, predicting cure from *in vitro* data is challenging and few work has been performed in this critical direction from a modelling perspective. Nevertheless, some efforts are underway using TKA data generated against *Mycobacterium ulcerans* to forecast the treatment shortening potential of novel drug combinations currently under clinical evaluation for Buruli ulcer treatment (*46, 48*).

### Limitations of the study

OPTIKA is essentially a TKA performed in a cost-effective high-throughput manner based on a surrogate fluorescence readout. Similar to static TKA, it presents common limitations: (*i*) Experiments described in this work have been performed with a standard laboratory strain from lineage 4 (*Mtb* H37Rv). Different strains including different lineages and clinical isolates should be compared to ensure genetically and drug susceptibility variability (*34*); (*ii*) We used standard *in vitro* growth conditions in synthetic 7H9 media containing glucose and glycerol as the main carbon sources, which promotes a relatively high homogeneity in the bacterial culture. However, the co-existence of different *Mtb* sub-populations at distinct physiological states with different drug susceptibility in patients is one of the reasons for the need of a multidrug regimen against TB (*49, 50*). Any combination of interest should be ideally further characterized under different experimental *in vitro* conditions mimicking these microenvironments, such as different carbon sources (i.e., glucose, cholesterol, fatty acids), low pH or non-replicating/starving metabolic states (*13, 25, 36, 51*). Although, the specific selection of *in vitro* assays to maximize translational output is still a matter of debate (*45*); (*iii*) Drug stability was not quantified in this study. To obtain a better understanding of the sterilizing capacity of the combinations, and specially its potential PAE, media stability of the different drugs in the combination should be performed over the time-course. This approach has been recently implemented to study *in vitro* the PK/PD relationships of the diarylquinoline TBAJ-587 and its metabolites against *Mtb*, suggesting that the real *in vitro* potency of these compounds was underestimated (*52*); (*iv*) OPTIKA should be considered as a high-throughput screening tool to evaluate large numbers of possible *n-*order drug combination and their potential lower order deconvoluted combinations and identify favorable (or not) drug partners. This is achieved by using a strategy of evaluating each drug in the combination at sub-optimal matching concentrations (i.e., 1/4xMIC or 1xMIC) with the rest of the drugs in the combo. These concentrations are microbiologically relevant (rooted on the MIC) but does not necessarily correlate with actual *in vivo* or clinical drug exposures, which could be too high and masked any potential interaction (as in our 4xMIC condition where drugs alone already exert all the measurable activity). Because OPTIKA’s activity readout is based on a fluorescence CFU-surrogate, integrating other biomarkers such as the ribosomal RNA synthesis (RS) ratio (*53*) could provide increase resolution. This biomarker could provide additional information on the effect of the drug combinations on the metabolic activity of the bacteria as a proxy for sterilization. Such approach has already been employed to study the performance of novel drug combinations for BU treatment (*54*); Finally, (*v*) in order to generate more *in vitro* relevant information, it is crucial to consider the dynamic drug concentration profiles of the drugs in the combination (*19*). In OPTIKA (and in static PK/PD systems in general such as TKA), drugs are added at a fixed concentration at time zero and they cannot be modulated over the experimental course in contract to other dynamic PK/PD systems, such as the hollow fiber system for tuberculosis (HFS-TB). The HFS-TB represents a powerful tool for simulating desired PK profiles, it is the only *in vitro* methodology endorsed by the European Medicines Agency as a predictive translational tool (*55*) and recent standardization efforts are consolidating it as a promising NAM (*56–58*). The HFS-TB enables derivation of relationships among drug exposure, bacterial kill rates, frequency of resistance, and the exposure needed for resistance suppression for single drugs and drug combinations. The model can predict, clinically relevant metrics, including susceptibility breakpoints, target dosages, and optimal drug combinations to understand and optimize the performance of potential anti-TB regimens through *in silico* modelling and simulation. However, the HFS-TB is costly and has a limited throughput capacity, which implies that drugs (and drug combinations) must be carefully selected to optimize resources and efforts dedicated to this assay. In this context, OPTIKA has the potential to accurately inform combinations to be tested in the HFS-TB model, a pipeline strategy already implemented in the ERA4TB Consortium (https://era4tb.org/).

## MATERIALS AND METHODS

### Bacterial strain, growth conditions and reagents

The *Mycobacterium tuberculosis* (*Mtb*) H37Rv (GenBank ID: NC_000962.3) strain was used for all experiments in this study. *Mtb* cells were routinely propagated at 37°C in Middlebrook 7H9 broth (Difco) supplemented with 10% Middlebrook albumin-dextrose-catalase (ADC, Difco), 0.2% glycerol and 0.05% (vol/vol) tyloxapol, or on Middlebrook 7H10 agar plates (Difco) supplemented with 10% (vol/vol) oleic acid-albumin-dextrose-catalase (OADC, Difco). An exponentially growing *Mtb* liquid culture (OD_600_ = 0.2-0.5) was aliquoted, stored at-80°C and frozen stocks CFU enumerated after one week. To enhance robustness of the assay, every experiment was started from a new frozen aliquot. Drugs used in this study were provided by GSK.

### Drug interaction assays based on growth inhibition

The day before of bacterial inoculation, tested compounds were dispensed in an 8×8 grid checkerboard format onto clear, flat-bottom 96-well plates (Costar 3599) in two-fold steps using an HP D3000 Digital Dispenser and stored at - 80°C: stock solutions of compounds were always prepared fresh on the same day of plate preparation. Plates were inoculated with *Mtb* cells (200 µL/well, 10^5^ CFU/mL) and incubated at 37°C. Internal controls included no drug (100% growth control) and no cells (0% growth control). After 6 days, 30 µL/well of MTT (2-(3,5-diphenyltetrazol-2-ium-2-yl)-4,5-dimethyl-1,3-thiazole bromide) at 5 mg/mL in Milli-Q water containing 20% (vol/vol) of Tween 80 (Sigma-Aldrich) were added to the assay plates and further incubated at 37°C for 24 hours. Plates were then equilibrated at room temperature for 30 min and adhesive seals placed on top. Absorbance at 580 nm (OD_580_) was measured with an EnVision Multilabel Plate reader (Perkin Elmer) and raw data normalized using positive and negative growth controls. Two analytical approaches were used to determine the degree of *in vitro* drug interactions against *Mtb*, which depended on the methodology used:

*(a) Checkerboard assay*: the lowest concentration of each single drugs that inhibited bacterial growth by 90% or 50% were selected as the IC_90_ or IC_50_ values, respectively. In order to quantify the degree of pairwise drug interaction, the Fractional Inhibitory Concentration Index (FICI) for a specific growth inhibition percentage was used and calculated as follow:

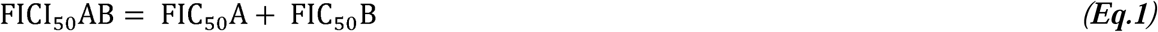

where the Fractional Inhibitory Concentration for a specific growth inhibition percentage was calculated as:

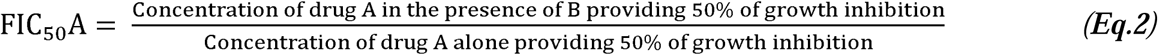

The FICI_90_ was similarly calculated using IC_90_ values. For a certain growth inhibition percentage (%), a synergistic interaction was defined by a FICI_%_ ≤ 0.5, antagonism by a FICI_%_ > 4 and, no interaction by FICI_%_ values from 0.5 to 4.

*(b) DiaMOND*: the % of growth of 2-fold serial dilutions for the single drugs and that of the equipotent combination mixture (diagonal of the 8×8 checkerboard layout) was analyzed using two approaches: (*i*) a dose response curve fitting analysis where the dose response of the single drugs and the dose response of the equipotent mixture were fitted to a four-parameter logistic model by the excel add-in XL fit version 5.5.0.5 (IDBS). IC_90_ and IC_50_ values were interpolated in the fitted curve equation. In case IC_90_ or IC_50_ values were higher than the highest concentration assayed, this was manually replaced by the next 2-fold increasing value of the highest concentration tested. (*ii*) a monotonically decrease analysis, where the % of growth was converted to a monotonically decreasing curve and the IC_90_ or IC_50_ values of single drugs and combinations, was interpolated from the corresponding linear drug concentration interval equation. FICI and FIC values were calculated using equations (*1*) and (*2*). Drug interactions were classified as described in Cokol *et al.* (*12*), *i.e.,* synergy, FICI_%_ < 0.85; antagonism, FICI_%_ > 1.1; and no interaction, 0.85 ≤ FICI_%_ ≤ 1.1.

### Minimum Bactericidal Concentration (MBC) assays

Bactericidal assays were coupled to growth inhibition assays as previously described (*25, 59*). Briefly, growth inhibition assay plates were inoculated with bacterial cultures and incubated at 37°C for six days. Prior to MTT addition, 20 µL/well were transferred to CARA plates. Inoculated CARA plates were placed in plastic bags and incubated at 37°C for nine days. Then, 40 µL/well of resazurin solution were added to the CARA plate, placed in plastic bags, and further incubated at 37°C for 24 hours. Plates were then equilibrated to room temperature for 30 min, sealed with adhesive films (Perkin Elmer, 6050185) and the top fluorescence (λ_exc_=530 nm; λ_emis_=590 nm) read with an EnVision Multilabel plate reader. Fluorescence background was calculated using wells containing media without inoculum. The MBC_99.9_ was calculated as the lowest drug concentration showing a fluorescence signal below the background. Similar to FICI calculations, the Fractional Bactericidal Concentration index was calculated as:

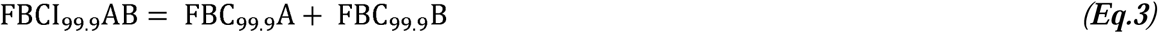

where for the single compounds:

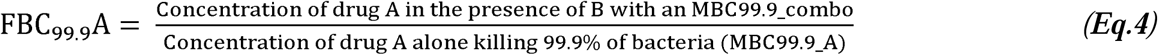

### Standard Time Kill Assays (TKA)

Frozen stocks of *Mtb* cells were thawed in 7H9/ADC medium to a density of *ca.* 10^4^ CFU/mL and incubated for 3 days (Day-3) to allow for bacterial recovery and exponential growth. At Day 0, this pre-inoculum (*ca*. 10^5^ CFU/mL) was evenly distributed (10 mL) in 25 cm^2^ tissue culture flasks and freshly prepared drug solutions added to the cultures at designated concentrations. At every time point, cultures were homogenized, aliquots (50 µL) 10-fold serially diluted in 1xPBS/0.1% tyloxapol up to the 10^-4^ or 10^-6^ dilution and plated (50/100 µL in duplicate) in 7H10/OADC quad petri dishes. Agar plates were incubated at 37°C for 2-3 weeks and colonies enumerated. Plates were checked again after 3-4 weeks of incubation to account for late growers. Cell density was reported as Log_10_CFU/mL.

### Optimized Time Kill Assays

CARA 96-well plates (Costar 3599) were prepared using Middlebrook 7H10 agar supplemented with 0.5% glycerol, 10% (vol/vol) OADC, 0.4% (weight/vol) activated charcoal (Sigma) and 0.5% (vol/vol) Tween 80, as described in Gold, B. *et al.* (*60*). Compounds were dispensed in Nunc™ Edge™ 96-well plates (Thermo Scientific) using the HP D300e digital dispenser. Combination assays were performed using MIC_90_ as the reference value. The plate layout included three test concentrations at matching concentrations, in technical quadruplicates: 1/4xMIC, 1xMIC and 4xMIC and compared to the activity of their respective single drugs.

*(i) OPTIKA calibration curve.* A *Mtb* culture with a known cell density (OD_600_= 0.125 equals to ca. 1×10^7^ cells/mL) was 10-fold serially diluted in 1xPBS/tyloxapol 0.1% and eight replicates of 20 µL/well plated onto CARA plates. The actual number of bacterial cells used for the calibration curves was determined by CFU enumeration.

*(ii) OPTIKA mother plates.* Plates with pre-dispensed compounds were inoculated with 250 µL/well of pre-inoculum culture (prepared as described for standard time-kill assays) and incubated at 37°C. At day 0 and every 7-10 days, 0.7-1 mL of sterile water were added to the plate edges to prevent from well evaporation. At every time-point, 20 µL/well from the mother plates were transferred to a CARA plate, placed in plastic bags and incubated at 37°C for 9 days. Then, a resazurin solution was added (40 µL/well) and plates were further incubated at 37°C for 24 hours in sealed plastic bags before fluorescence measurement in an EnVision Multilabel plate reader (λ_ex_= 530 nm/λ_em_= 590 nm, top read).

### Statistical analysis of OPTIKA and combinations classification

A calibration curve from a culture with known cell density was generated for every time-point of the kill kinetic, as above described. For every cell density, the fluorescence mean +/-SD of 8 replicates was plotted versus the actual Log_10_CFU/mL. At every time point, fluorescence background and lower linear range of the calibration curve defined the limit of detection for sample interpolation. The linear range of the calibration curve was fitted to a linear regression and the equation was used to calculate the bacterial concentration in the samples. Relative Fluorescence Units (RFU) of sample plates were converted to Log_10_CFU/mL through data interpolation into the calibration curve equation of the corresponding time-point. When the observed value was below the lower limit of detection of the corresponding calibration curve, this was manually corrected to the limit of detection. To classify the degree of interaction of the combinations, the mean of Log_10_CFU/mL of every drug condition at every time-point was calculated with GraphPad Prism software. Five parameters were studied:

i. *Bacterial killing rate.* Calculated as the mean Log_10_CFU/mL at day 2 (D2) minus the mean Log_10_CFU/mL at day 0 (D0), divided by 2. Graphically, this corresponded to the slope of the killing curve from day 0 to day 2 *(**Eq.5**)*. This value was normalized by subtracting the slope of the untreated control, calculated with *(**Eq.6**)* and lead to the calculated parameter of the normalized slope to untreated *(**Eq.7**)*.

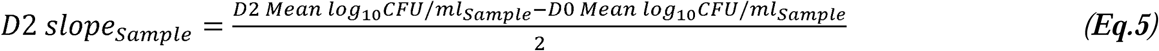

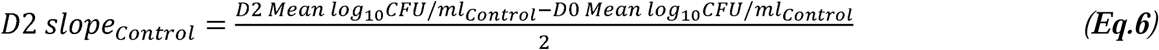

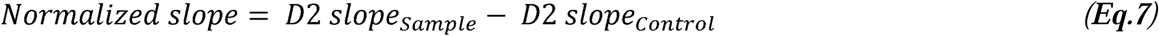

In order to determine whether combinations killed faster than the most active drug alone, normalized slope values of the combinations were compared with those of the respective single drugs. Based on this approximation, combinations were classified as: Favorable Interaction (FI), if this parameter was significantly lower for the combination than for the most active single drug; Non Favorable Interaction (NFI), if this parameter was significantly higher for the combination than for the most active single drug; No Interaction (NI), if difference between the combination and the most active drug alone were not significantly different and; Not determined (ND), if the limit of detection was reached at day 2 by at least, one of the single drugs.

ii. *Bacterial burden reduction at Day 4 (D4):* the bacterial burden of the combinations was compared with that of the related single drugs at day four of the kill kinetic assay. Considering the generic pair-wise drug combination constituted by A and B, log reduction of single drugs was calculated as follows:

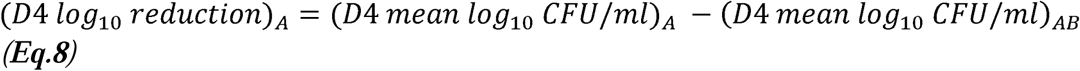

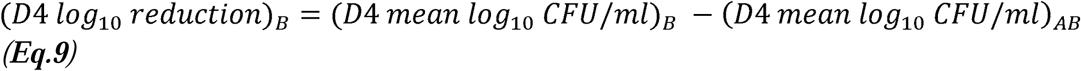

The minimum absolute value of log reduction of A and B was selected to calculate the D4 log reduction of the combination, using the formula:

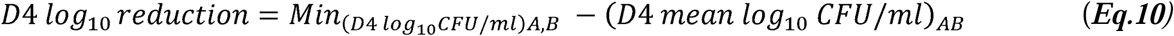

Combinations were then classified according to the D4 Log_10_ reduction parameter as: Favorable Interaction (FI), if the reduction was equal or higher than 2-Log_10_CFU/mL compared to effect of the most active drug alone in the combination; Non Favorable Interaction (NFI), if there was an increase equal or higher than 2-Log_10_CFU/mL compared to effect of the most active drug alone in the combination; No Interaction (NI), if differences between the combination and the most active drug alone were within a 2-Log_10_CFU/mL range and; Not determined (ND), if the limit of detection was reached at day 4 by, at least, one of the single drugs.

iii. *Bacterial burden reduction at D7*. This parameter was calculated as described for D4 Log_10_ reduction, but with data from D7.

iv. *Regrowth at D21:* for single drugs and combinations, regrowth was a categorical parameter, indicating whether the bacterial load was above or within the limit of detection (LD). Calculated as follows:

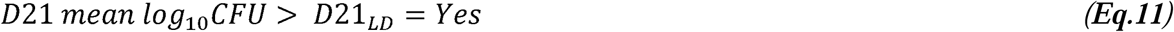

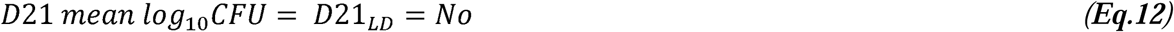

By comparison of the ability of combinations and related single drugs to prevent regrowth, combinations were classified as follows: Favorable Interaction (FI) was attributed to combinations with no regrowth (**Eq.12**), compared to the corresponding single drugs when these were not able to prevent bacterial regrowth (**Eq.11**); No Interaction (NI), was defined when both the combination and single drugs in the combination displayed bacterial regrowth; Non-Favorable Interaction (NFI) was defined in combinations that showed bacterial regrowth (**Eq.11**), whereas any of the single drugs bacterial load was at the limit of detection (**Eq.12**); Not determined (ND), if regrowth was not observed in, at least, one of the single drugs.

v. *Regrowth at endpoint:* this parameter was calculated as described for D21 regrowth, but for the last time-point of the kill kinetic (*ca*. D49, 7 weeks of incubation).

All analyses were performed using GraphPad Prism version 8.1.1 (La Jolla, CA). Mann-Whitney U test was used to compare two groups and Kruskal-Wallis test (followed by Dunn’s multiple comparison test) was used to compare parameters among three or more groups. All analyses were considered statistically significant at P value <0.05.

## ACKNOWLEDGMENTS

We would like to thank the TCOLF management team, especially Félix Calderón, for their support carrying out this work. We would like to acknowledge Carlos Martín for the kind gift of the *M. tuberculosis* H37Rv strain used in this study, and Jordana Galizia for critical input discussion and for providing Figure S1, both from the University of Zaragoza.

## Author contributions

CRediT (Contributor Roles Taxonomy) has been applied for author contribution. *Conceptualization:* MPAC, AML, SFB, RGdR and SRG; *Data curation:* MPAC and PG; *Formal Analysis:* MPAC and PG; *Funding acquisition:* SRG; *Investigation:* MPAC; *Methodology:* MPAC and SRG; *Project administration:* SFG, RGdR and SRG; *Resources:* PG; *Software:* PG; *Supervision:* AML, SFB and SRG; *Validation:* MPAC and PG; *Visualization:* MPAC and SRG; *Writing – original draft:* MPAC, AML, SFB and SRG; *Writing – review & editing:* MPAC, AML, SFB and SRG.

## Data and materials availability

All data pertaining to this work is within the main manuscript or supplementary information. Primary data are available from the corresponding author upon request.

