## Supplementary Tables and Figures for "OPTIKA, a new high content kill-kinetic assay to longitudinally assess *in vitro* drug combinations against *Mycobacterium tuberculosis*"

***Running title:*** OPTIKA, a new high content kill-kinetic assay

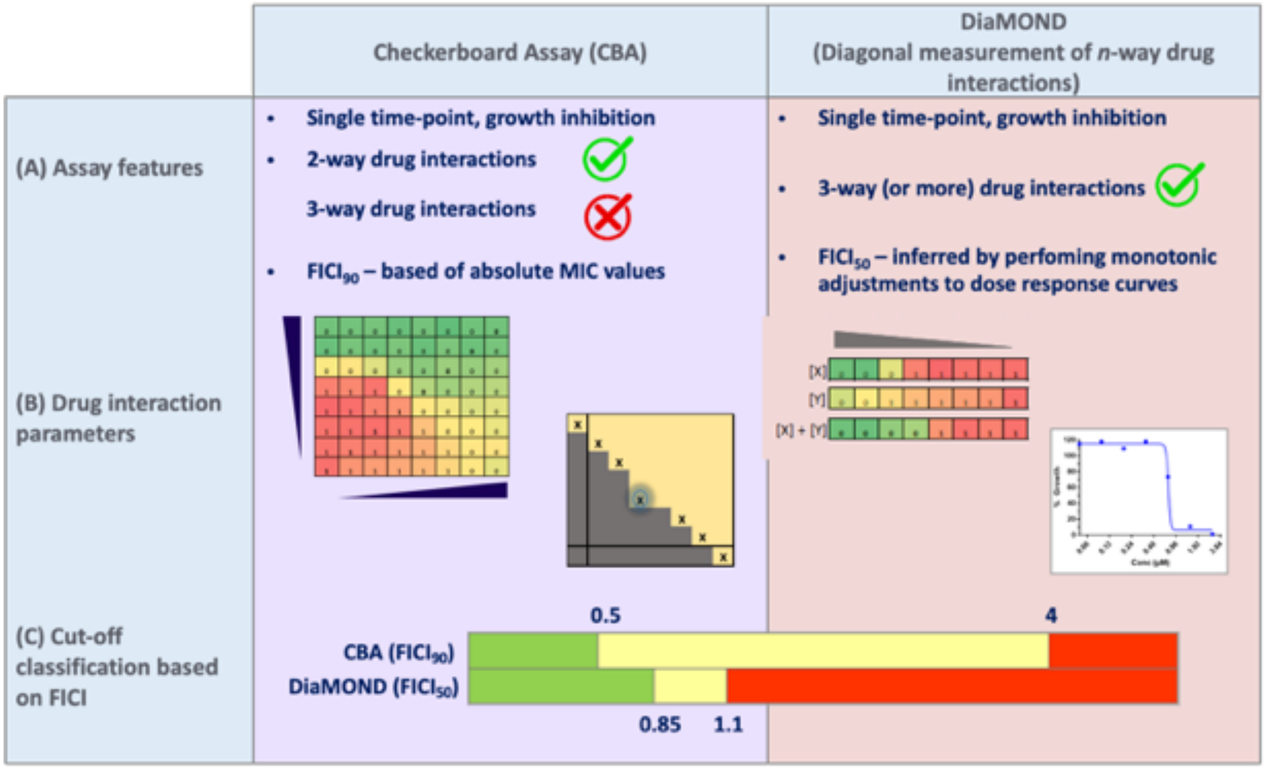


**Figure S1. Comparison between the checkeboard and DiaMOND assays. (A)** Both methodologies are based on growth inhibition assays. DiaMOND provides the capacity to interrogate *n*-way drug combinations, while CBA is limited to 2-way drug combinations (up to 3-way could be performed but at the cost of intense experimental performance). **(B)** DiaMOND simplified the CBA by only testing the dose response of the drugs alone and the equipotent mixture of the two compounds. Calculations of the Fractional Inhibitory Concentration Index (FICI) differ between CBA and DiaMOND. While CBA calculate $FICI_{90}$ based on absolute MIC values, DiaMOND calculate $FICI_{50}$ based on $IC_{50}$ calculations from monotonically adjustments of the dose response curves. **(C)** The actual FICI values used for categorization of the drug combinations differ in both CBA and DiaMOND. DiaMOND overrepresents antagonistic interactions while CBA classifies most of the combinations as No Interaction.

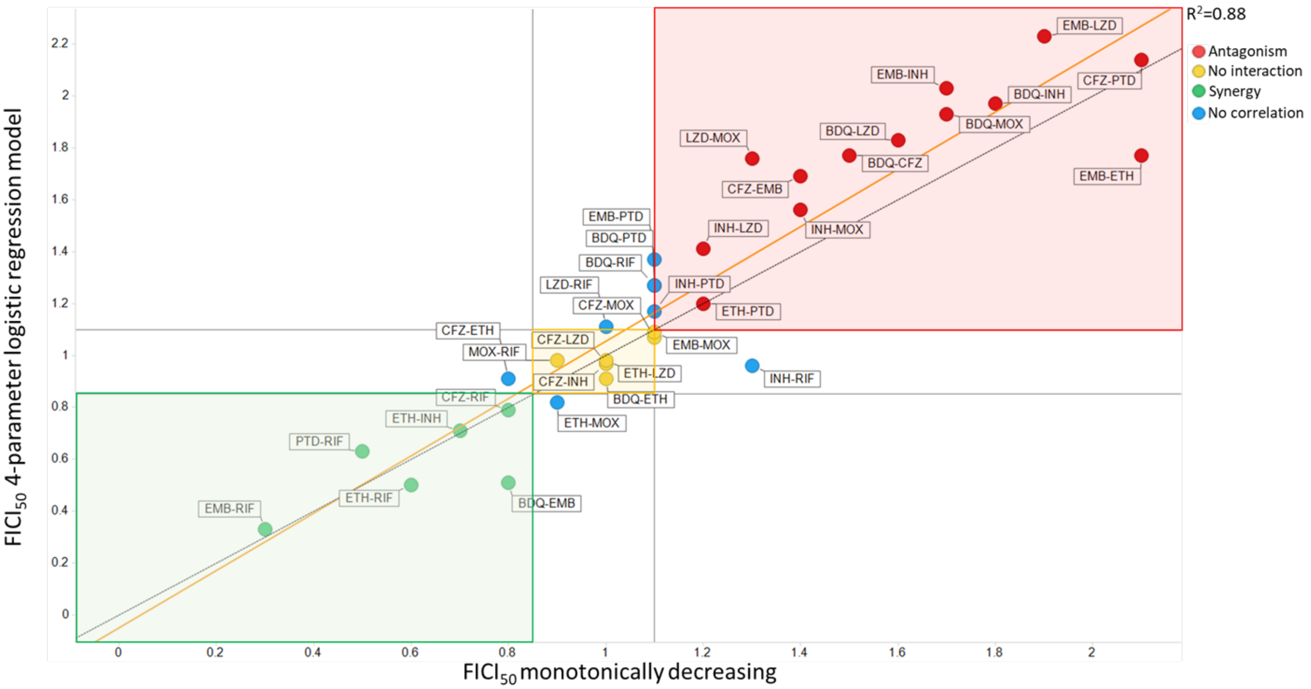


**Figure S2. Drug interaction classifications based on the DiaMOND methodology according to the method of analysis.** A four-parameter logistic regression model is compared to a monotonical decrease method for calculations of the $FICI_{50}$ of each pairwise combination. Circle legends: green: synergy. Yellow: no interaction. Red: antagonism. Blue: no matching combinations. Background zone legends; green: area for synergy combinations classified by both methods; yellow: area for no interaction combinations; white: area for combinations classified differently by both methods. Straight dotted line: perfect correlation zone (x=y). Orange line: correlation line between two sets of results ($R^2$ = 0.88). Bedaquiline (BDQ), clofazimine (CFZ), ethambutol (EMB), ethionamide (ETH), isoniazid (INH), linezolid (LZD), moxifloxacin (MOX), pretomanid (PTD), rifampicin (RIF).

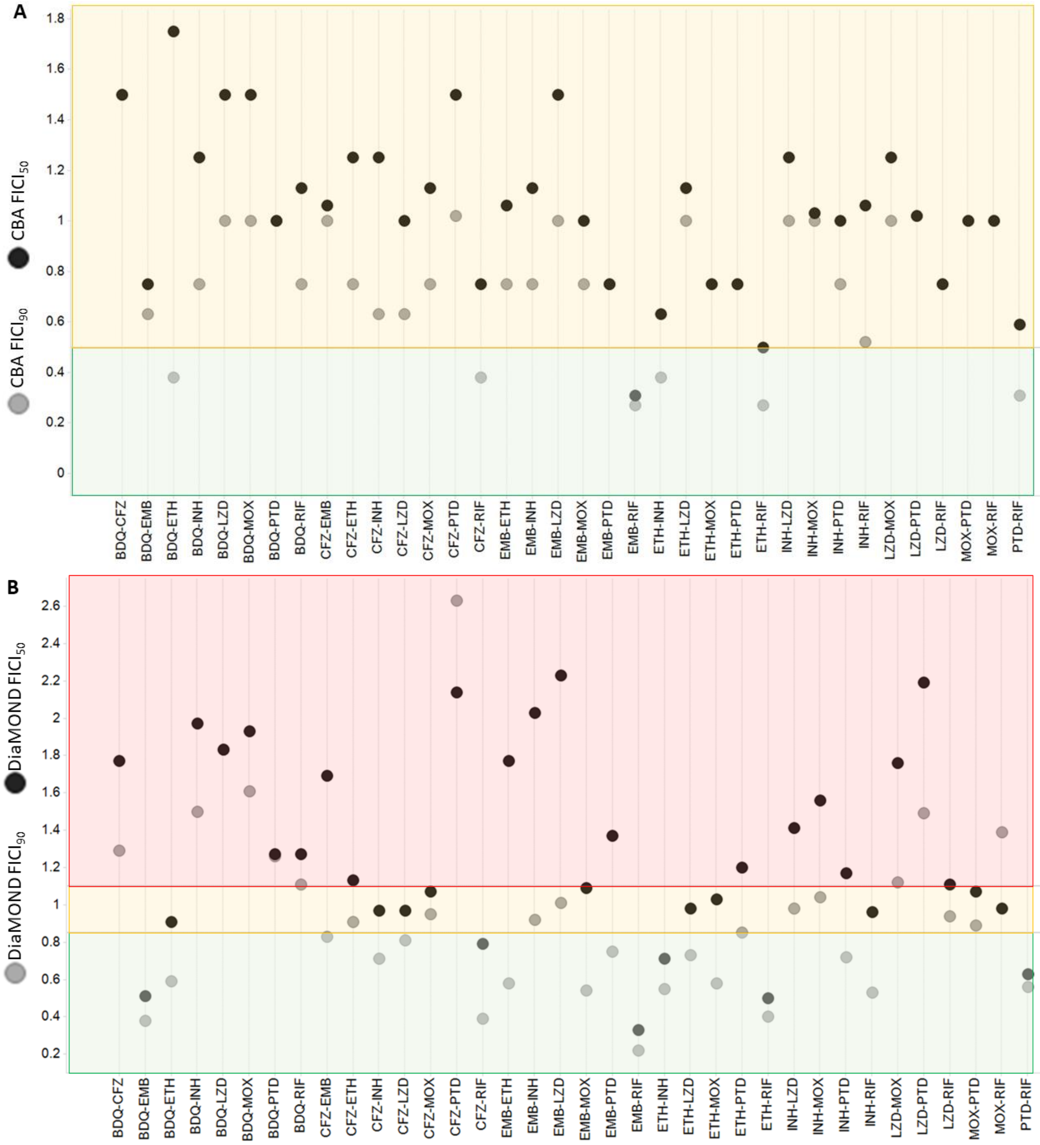


**Figure S3. FICI$_{50}$ *vs.* FICI$_{90}$ comparison.** Data generated in this work for the 36 pair-wise combinations was analyzed based on both IC$_{50}$ and IC$_{90}$ values and according to CBA **(A)** and DiaMOND **(B)** methodologies. Grey circles: FICI$_{90}$. Black circles: FICI$_{50}$. Green background: synergy. Yellow background: no interaction. Red background: antagonism. Bedaquiline (BDQ), clofazimine (CFZ), ethambutol (EMB), ethionamide (ETH), isoniazid (INH), linezolid (LZD), moxifloxacin (MOX), pretomanid (PTD), rifampicin (RIF).

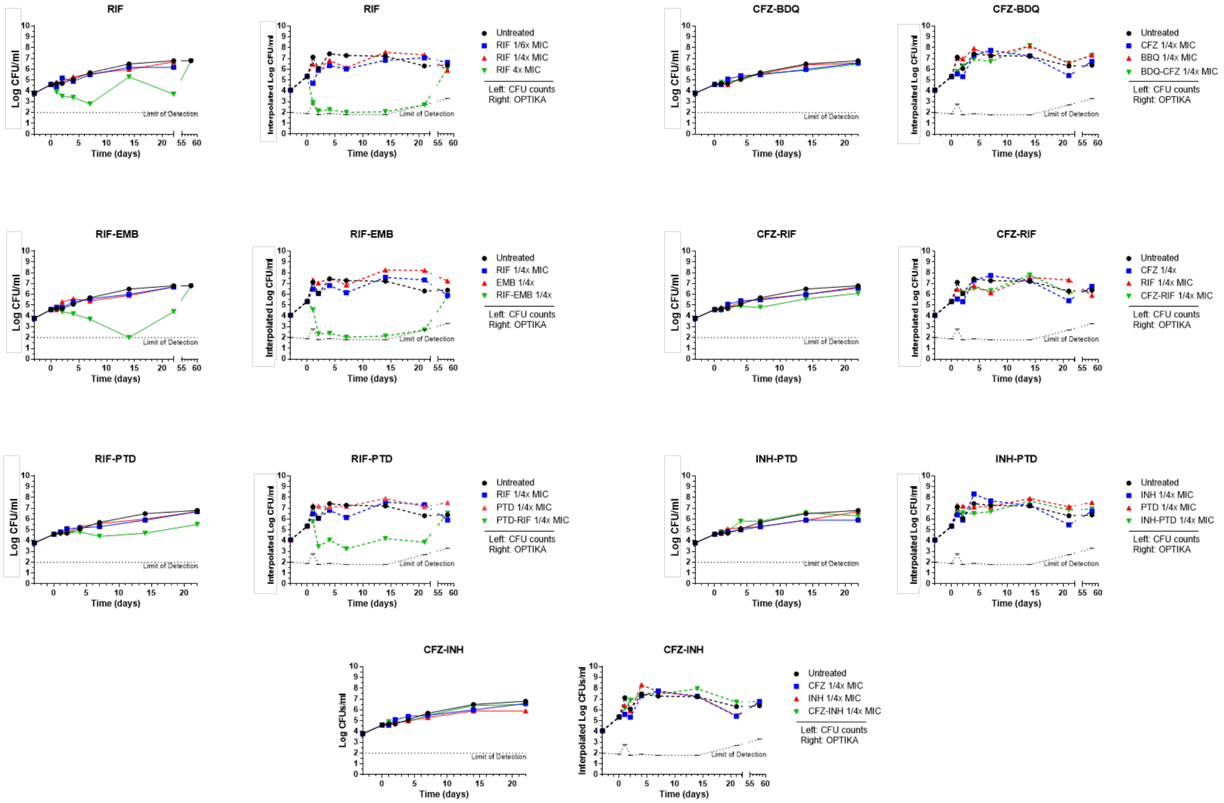


**Figure S4. Comparison of traditional TKA versus OPTIKA.** Head-to-head comparison of time-kill kinetic determinations measured by traditional CFU-based TKA and the fluorescence-based OPTIKA methodology. **(A)** Drugs and MIC values used. **(B to H)** Killing curve profiles generated by traditional TKA (left plot of every pair) and OPTIKA (right plot). Concentrations used relate to the individual MIC values of each drug were: Bedaquiline, BDQ, 0.07 µM; Clofazimine, CFZ, 0.38µM; Ethambutol, EMB, 12.97 µM; Isoniazid, INH, 3.52 µM; Pretomanid, PTD, 0.62µM; and Rifampicin, RIF, 0.40 µM.

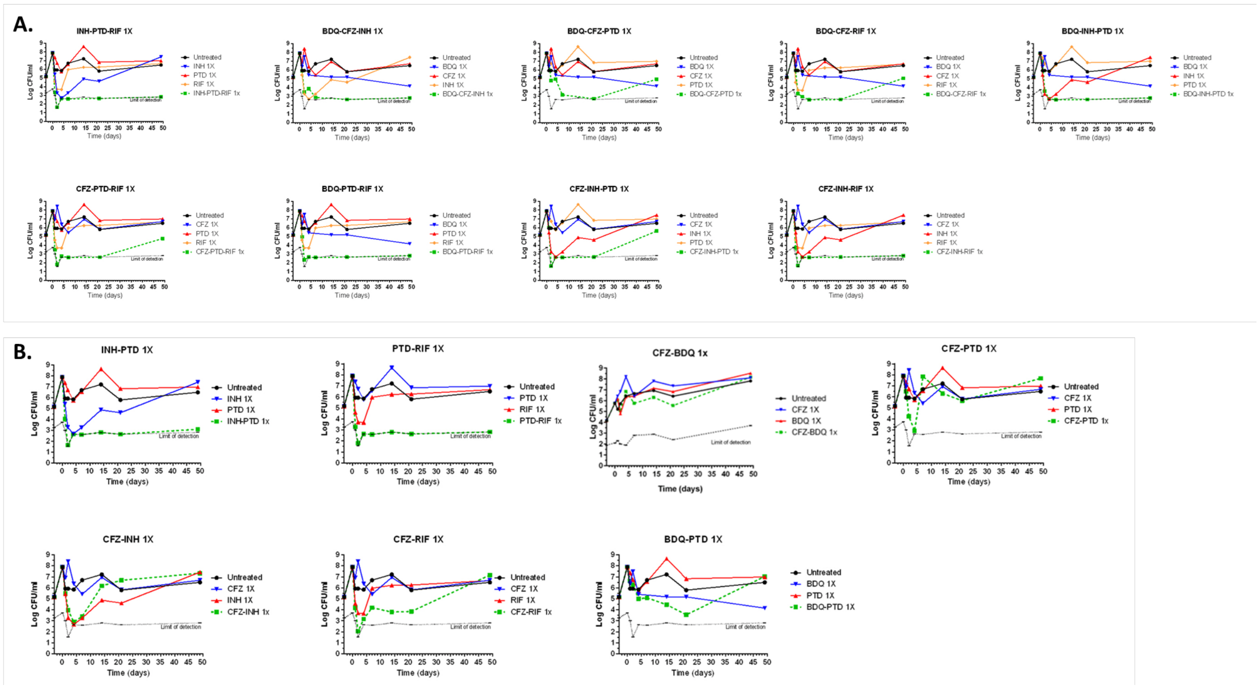


**Fig. S5. OPTIKA triple combinations.** Killing curve profiling of the triple combinations at 1x MIC shown in Fig. 6. **(A)** Triple combination and respective single drugs. **(B)** Pair-wise combinations. Bedaquiline (BDQ); Clofazimine (CFZ); Isoniazid (INH); Pretomanid (PTD); Rifampicin (RIF).

**Table S1. Performance comparison of OPTIKA versus traditional time-kill assays.** OPTIKA allows for a ca. 1,000-fold increase in sample processing by an individual operator and increase data robustness since quadruplicate biological samples are tested for an estimated total of 770 unique conditions. Because OPTIKA is based on the CARA assays, which measures fluorescence resazurin conversion, time to readout can be shortened to 10 days from the typical 2-4 weeks needed until CFU are visible. Working in a 96-well plate format allows the use of multichannel pipettes instead of handling cultures in flask, thus reducing manipulation times in the BSL3. TKA, Time Kill Assay. OPTIKA, Optimized Time Kill Assay.

| | **Traditional TKA** | **OPTIKA** |
|---|---|---|
| **Volume of the sample** | 10 mL (flask) | 250 µL (96-well plates) |
| **Handling** | Serial dilutions | Multichannel handling |
| | 2x Technical duplicates | 4x Biological replicates |
| | Culture agar plating | CARA assay |
| | CFU enumeration | Fluorescence reading |
| | 4-6 hours per time point | 3-4 hours per time point |
| **Throughput** | Max. 30 samples/operator | 88 samples/plate (up to 35 plates): 3080 samples/operator |
| | 30 unique conditions | 770 unique conditions in technical quadruplicates |
| **Time to readout** | ca. 3 weeks | 10 days |
